# GABAergic inhibition in the human visual cortex relates to eye dominance

**DOI:** 10.1101/2020.09.10.291047

**Authors:** I. Betina Ip, Claudia Lunghi, Uzay E. Emir, Andrew J. Parker, Holly Bridge

## Abstract

Our binocular world is seamlessly assembled from two retinal images that remain segregated until the cerebral cortex. Despite the coherence of this input, there is often an imbalance between the strength of these connections in the brain. ‘Eye dominance’ provides a measure of the perceptual dominance of one eye over the other. Theoretical models suggest that eye dominance is related to reciprocal inhibition between monocular units in the primary visual cortex, the first location where the binocular input is combined. As the specific inhibitory interactions in the binocular visual system critically depend on the presence of visual input, we sought to test the role of inhibition by measuring the concentrations of inhibitory (GABA) neurotransmitters during monocular visual stimulation of the dominant and the non-dominant eye. GABA-levels were acquired in V1 using a combined functional magnetic resonance imaging (fMRI) and magnetic resonance spectroscopy (MRS) sequence on a 7-Tesla MRI scanner. Individuals with stronger eye dominance had a greater difference in GABAergic inhibition between the eyes. This relationship was present only when the visual system was actively processing sensory input and was not present at rest. We provide the first evidence that imbalances in GABA levels during ongoing sensory processing are related to eye dominance in the human visual cortex. This provides strong support to the view that intracortical inhibition underlies normal eye dominance.

**SIGNIFICANCE STATEMENT:** What we see is shaped by excitation and inhibition in our brain. We investigated how eye dominance, the perceptual preference of one eye’s input over the other, is related to levels of inhibitory neurotransmitter GABA during monocular visual stimulation. GABAergic inhibition is related to eye dominance, but only when the visual system is actively processing sensory input. This provides key support for the view that imbalances in visual competition that are observed in the normal visual system arise from an inability of GABA signalling to suppress the stronger sensory representation.

## INTRODUCTION

Eye dominance in the healthy visual system is the preference for using one eye over the other in a visual task [1]. Extreme eye dominance is associated with amblyopia, a neurodevelopmental disorder that causes a chronic loss of normal monocular and binocular function with an incidence of 2-4% in the general population [2-4]. Input from the two eyes is combined for the first time in the primary visual cortex to serve binocular vision. Therefore processing at this stage is thought to be decisive in eye preference. Thus, understanding the relationship between eye dominance and the brain is an opportunity to investigate the neural mechanisms that the cerebral cortex selectively uses to serve perception.

Evidence for a neural mechanism of eye dominance comes from the abnormal binocular visual system. Children who grow up with one deviating eye often develop strabismic amblyopia. In typical adult life, double vision results when the two eyes receive inputs that do not correspond. This is associated with severe discomfort and headaches. Yet many strabismic amblyopes perceive only a single image, not two. This is because their visual system has prevented double images from reaching perception by making the input from the deviating eye non-visible and relaying the image from the non-deviating eye. This process is called binocular suppression. Chronic suppression could be ‘nature’s way out of trouble’ [5]. The drawback of suppression is that it is related to a loss of binocular visual functions [6], most notably stereopsis and binocular summation. Hence, visual cortex suppression has been named as one of the prime causes of the perceptual deficits observed in amblyopia [7].

Severely imbalanced vision causes anatomical abnormalities in the primary visual cortex [8], with functional abnormalities likely maintained by GABAergic inhibition [9]. The strongest evidence in support of this view has come from pharmacological studies in amblyopic animal models. In strabismic amblyopic cats, localized application of the GABA_A_ receptor antagonist, bicuculline, to V1 reduced binocular suppression [10]. In amblyopic rats, pharmacologically decreasing GABAergic signaling in the visual cortex has been linked to a recovery of normal structure and visual function in the amblyopic eye [11, 12]. This recovery was blocked by cortical delivery of diazepam, a GABA_A_ agonist [11]. These studies provide strong support for a role of GABAergic inhibition in maintaining extreme eye dominance in the adult brain.

Amblyopic suppression may be an extreme form of normal binocular interaction, revealed in a subtle form in normal participants using binocular rivalry [9]. Binocular rivalry is a standard method to quantify sensory eye dominance in the normal-sighted population [1]. It induces continuous perceptual alternations between incongruent images presented separately to the left or right eye [13, 14].

Theoretical views suggest that intracortical inhibition could explain the pattern of perceptual dynamics [15, 16]. In recent years, support for this view has come from studies applying magnetic resonance spectroscopy (MRS) to measure GABAergic inhibition in the human brain. These reports found that binocular rivalry dynamics in healthy participants are correlated to and caused by baseline GABA levels in the early visual cortex [17, 18] [19]. In addition, transient increases in eye dominance duration have been correlated with lower GABA levels in the primary visual cortex [20]. These results have highlighted the role of baseline GABAergic inhibition in binocular rivalry, yet the hypothesis that imbalances during active stimulation of each eye are related to eye dominance awaits critical support.

To test the long-standing prediction that imbalances in cortical inhibition during monocular visual response are related to eye dominance [9], we set out to measure GABA during functional stimulation. Specifically, we wanted to quantify the ‘interocular’ difference in GABA during monocular stimulation of either the stronger or the weaker eye. Conventionally, MRS experiments measure GABA levels during the resting state, when participants have their eyes closed, or during an active state in which subjects simply watch a movie with both eyes open. In absence of a specific task, such measurements at rest are thought to reflect the overall inhibitory tone in the brain area under study [21]. In contrast, our study takes advantage of recent advances in ultra-high field functional MRS, that allow measurement of metabolite levels during task performance [22-25]. For all participants, we determined both eye dominance using binocular rivalry and the magnitude of interocular inhibition, manifest in the difference in GABA levels during stimulation of the stronger and weaker eyes respectively. Our results provide the first evidence that eye dominance in the human visual cortex is related to neural mechanisms that deliver active interocular inhibition.

## RESULTS

For each participant, sensory eye dominance was measured individually using binocular rivalry stimuli presented by means of a Wheatstone stereoscope (Fig 1A). Binocular rivalry phase distributions pooled across all observers are not normally distributed (Shapiro-Wilk Normality Test, W-stats = 0.64, *p* = 0.0001) as demonstrated by the typical skewedness [26] of bistable percept phase durations (Fig. 1B). The median phase durations were used as a measure of central tendency [27]. Median phase durations across 12 observers were normally distributed (W-stats = 0.95, *p* = 0.337), hence parametric statistics used for significance testing, with an uncorrected alpha of 0.05. After confirming that there was no overall difference in right vs left eye median phase durations (two-tailed paired samples *t*-test, *t*(11) = 0.228, *p* = 0.824), eye dominance was assigned based on the length of the median phase duration (Fig 1C, left; 5 observers = left eye dominant, 7 observers = right eye dominant, left vs right median). Not surprisingly, median phase durations were greater for the DE compared to the NDE (two-tailed paired samples *t*-test, *t*(11) = 2.960, *p* = 0.013). To obtain a singular measure of the increase in eye dominance, an eye dominance index (Fig 1C right; ‘EDI’, see Methods Equation 1) was calculated. The EDI measures the increase in duration of the DE phase as a percentage of the NDE phase (range: 0.64 – 23.1%, mean: 7.9 ± 8.2%).

**Figure 1.**
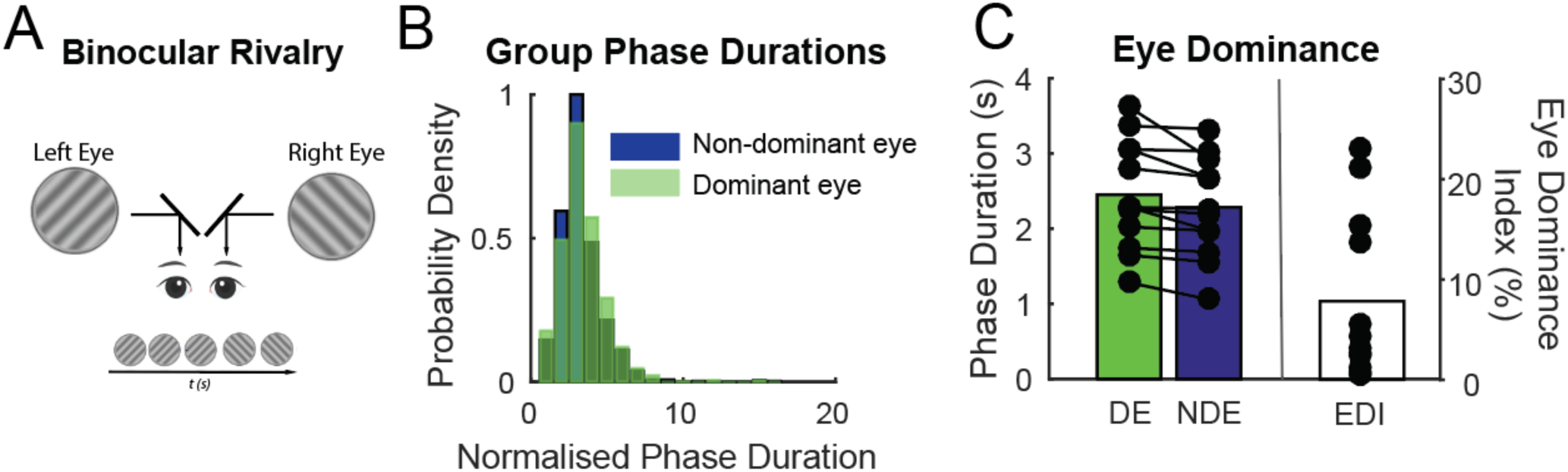
**(A)** Sensory eye dominance was measured in a binocular rivalry experiment outside of the scanner. Stimuli were presented on a mirror stereoscope and participants reported their subjective percept using a continuous key press. **(B)** Histograms of phase distributions for the dominant (DE, green) and non-dominant (NDE, blue) eye were pooled across participants. Data were normalised by the median dominant eye phase duration. **(C)** Average phase durations for DE (green) and NDE (blue) are plotted on the left. On the right are the biases in phase duration quantified for each participant by the ratio of the dominant eye median phase duration (s) / non-dominant eye phase duration (s). This index is henceforth referred to as ‘Eye Dominance Index’ (clear bar, EDI).

Mutual inhibition between neural inputs from each eye to the cortex is thought to drive eye dominance. To compare inhibition measured during alternating stimulation of the dominant and the non-dominant eyes, we calculate the ‘interocular GABA’ metric. The interocular GABA quantifies the difference in GABA levels between DE and NDE as a proportion of the NDE GABA concentration (see Fig 2A, also Methods, Equation 3). We then related eye dominance to the interocular GABA metric in early visual cortex. Fig 2B shows that EDI was significantly correlated with interocular GABA (Fig. 2B, GABA:H_2_0, Spearman’s Rank Correlation *r* = 0.61, uncorrected p= 0.037; GABA:tCr, r = 0.60, uncorrected p = 0.042). More detailed breakdown of this relationship shows that there is no correlation between EDI and GABA in the DE (Fig 2C: Spearman’s Rank Correlation, GABA:H_2_0, r = 0.01, p = 0.97; GABA:tCr, r = 0.02, p = 0.93), but a trend towards a relationship of EDI with GABA in the NDE (Fig 2D: Spearman’s Rank Correlation, GABA:H20, r = -0.44, p = 0.14; GABA:tCr, r = -0.48, p = 0.11). We conclude that individuals with stronger eye dominance also have a greater *difference* in GABAergic response between their dominant and non-dominant eyes. Across participants, GABA during DE compared to NDE viewing was close to statistical significance (GABA:H_2_0, paired t-test, t(11)= 2.09, p =0.06; GABA: tCr, paired t-test, *t*(11) = 1.75, *p* = 0.108). To evaluate whether visual stimulation was necessary to reveal the relationship, we also correlated EDI with resting GABA levels. There was no relationship between eye dominance and the level of GABA in the resting condition (GABA:H_2_0, r = - 0.129, p = 0.7; GABA:tCr, r = -0.09, p =0.76).

**Fig. 2.**
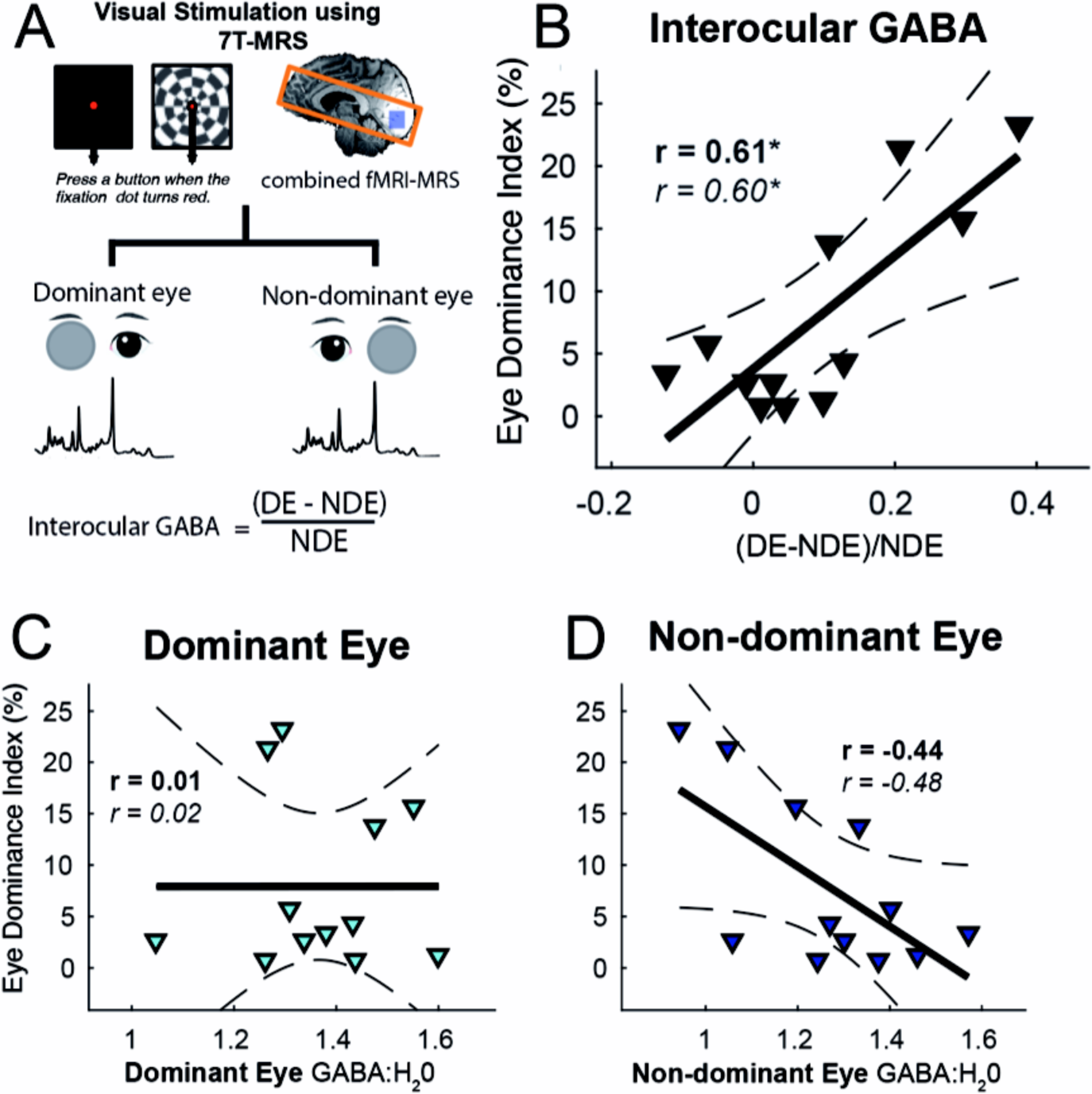
**(A)** Metabolite concentrations were collected using a monocular visual stimulation protocol in the 7-Tesla MRI scanner. Participants viewed visual stimuli with either the left or right eye, while the non-viewing eye was covered using a semi-translucent occluder. Conditions (‘left eye’, ‘right eye’) were collected in separate, pseudo-randomised runs balanced for order effects. During visual stimulation, participants performed a constant fixation task, while four alternations of a full-screen blank black screen (64 s) and a 50% contrast flashing checkerboard (64 s) were presented in the background. Data were collected using a combined fMRI-MRS sequence that acquired interleaved MRS and BOLD-EPI data. The diagram of a standard brain shows the field-of-view for EPI acquisition (red box) and the 2×2×2 cm MRS voxel within the occipital cortex (blue square). Interocular GABA is quantified as the difference between DE and NDE GABA, as a proportion of NDE GABA. **(B)** Eye dominance was highly correlated to the difference in GABA between DE and NDE eyes (‘interocular GABA’). **(C)** Eye dominance is uncorrelated with GABA:H_2_0 during stimulation of the DE. **(D)** Eye dominance is showing a trend towards correlation with GABA during stimulation of NDE. Solid line represents best fitting linear model, hatched lines represent 95% confidence interval of the regression line. r = Spearman’s Rank Correlation Coefficient, * = uncorrected p < 0.05. r-values in italics represent results for GABA scaled to total Creatine.

### Eye dominance was not correlated to difference in BOLD-signal to flashing checkerboard stimuli between DE and NDE eyes

Next, we investigated the relationship between eye dominance and the BOLD-fMRI response, which were acquired in the same time point and anatomical region as the MRS data (see Fig. 2A). While MRS values can be measured across the entire scan run, the measurement of the BOLD-signal requires a baseline comparison. Thus, the BOLD measurement reflects the comparison of viewing flashing checkerboard blocks relative to the fixation task with one eye while the other eye is occluded. In contrast, the MRS analyses compare viewing of both fixation task and the checkerboard blocks with one eye while the other eye is occluded. We found no significant difference in BOLD-signal between DE and NDE visual stimulation (Fig. 3A: Wilcoxon’s Rank Sum, z=-0.77; p = 0.43). We then calculated the interocular BOLD-signal (for ‘interocular BOLD’, see Methods Equation 3). There was no correlation between EDI and interocular BOLD (Fig. 3B, Spearman’s *r* = 0.251, *p* =0.430). We also tested whether the difference in the GABA response to the flashing checkerboard compared to fixation alone (ΔGABA) was related to eye dominance. We found no correlation between eye dominance and ΔGABA (GABA: H_2_0, r = 0.24, *p* = 0.44; GABA:tCr, r = 0.24, p =0.44), suggesting that neurochemical responses to flashing checkerboards are not systematically associated with eye dominance.

**Figure 3.**
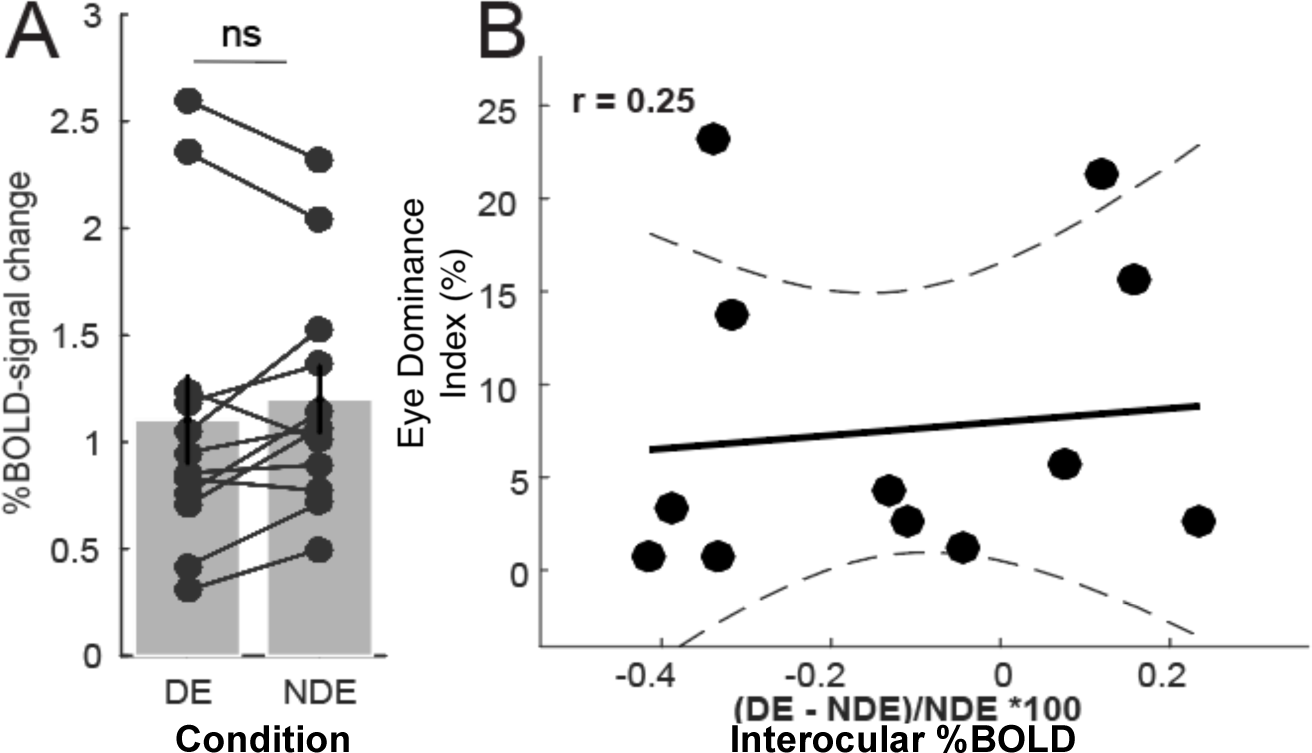
The BOLD-signal change evoked by flashing checkerboards during DE and NDE viewing. **(A)** Dominant and non-dominant eye BOLD changes did not show a difference. **(B)** There was no relationship between eye dominance and the interocular %BOLD. Errors are 1 ± SEM. r = Spearman’s Rank Correlation. Solid line represents best fitting linear model, hatched lines represent 95% confidence interval of the regression line. ns = not significant.

### Resting GABA relates to binocular rivalry suppression

A complementary measure of binocular rivalry dynamics, the proportion perceptual suppression, has been shown to correlate with resting GABA in the healthy brain [18]. For direct comparison with these findings, we calculated binocular rivalry suppression (BRS), a metric that takes into account the proportion of ‘mixed periods’ (Fig. 4A). Measures of BRS were normally distributed and Pearson’s linear correlation coefficient was used to relate perceptual suppression to resting GABA:tCr levels. We replicate the correlation between perceptual suppression and resting GABA:tCr (Fig. 4B, r = 0.64, p = 0.025). However, the correlation was not significant when metabolites were scaled to water, indicating possible influences from data quality.

**Figure 4.**
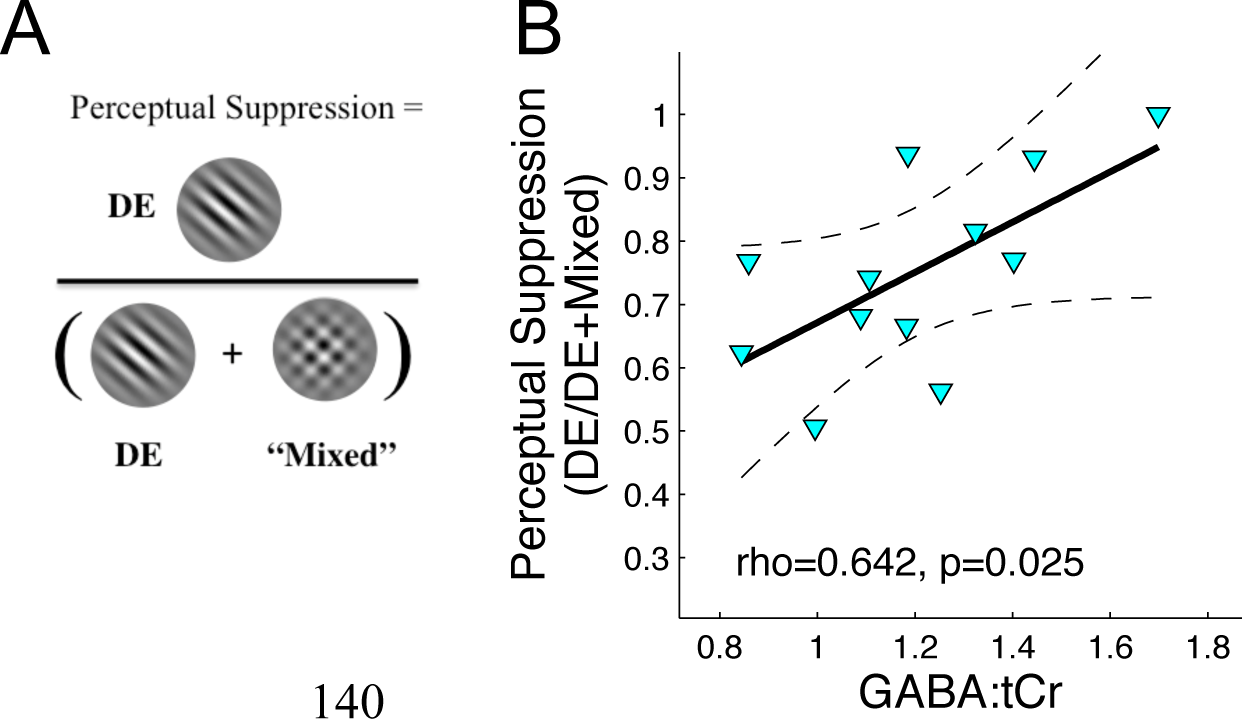
To compare with previous data, we calculated an alternative measure of binocular rivalry dynamics, the ‘perceptual suppression’ rate (**A**; Robertson et al., 2016). **(B)** Resting GABA:tCr levels correlated to binocular rivalry perceptual suppression [dominant eye / (dominant eye + mixed percept)]. r = Pearson’s linear correlation co-efficient r. Solid line represents best fitting linear model, hatched lines represent 95% confidence interval of the regression line.

### General quality differences and BOLD effects did not influence GABA estimates during visual stimulation

To investigate whether spectral quality and BOLD effects in the MRS-spectra influenced GABA estimates, we pooled GABA values across visual stimulation conditions and correlated them with spectral SNR. We calculated SNR as the height of the tNAA peak divided by the standard deviation of a region of noise, and correlated the SNR to GABA and found no relationship (Fig. 5A, GABA:H_2_0, Pearson’s Linear Correlation, r = 0.02, p = 0.89; GABA:tCr, Pearson’s Linear Correlation, r = 0.18, p = 0.381).

**Fig. 5:**
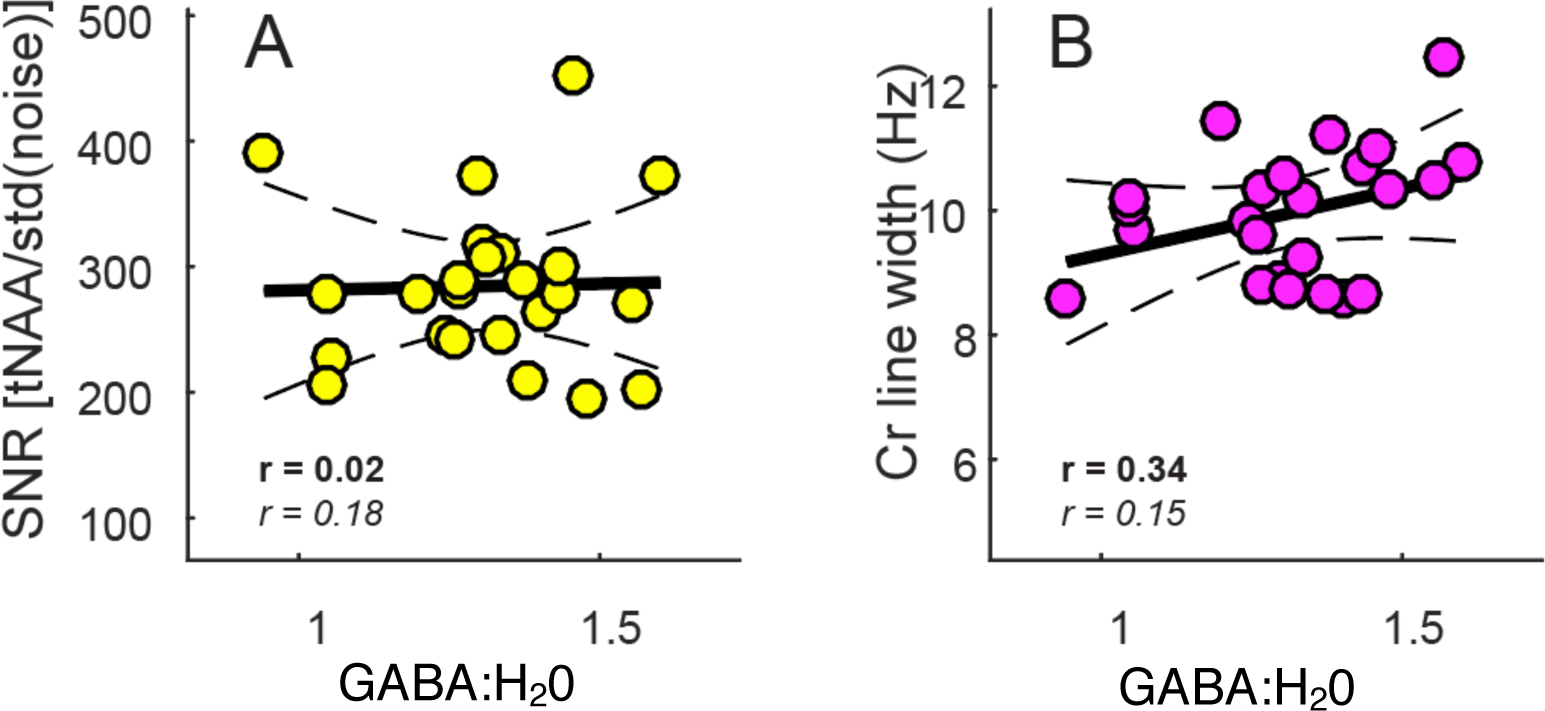
Data quality measures and GABA estimates. **(A)** Pooled GABA:H_2_0 values across viewing conditions and correlation to SNR. **(B)** GABA correlation to tCR LW. r = Pearson’s Linear Correlation Co-efficient. Solid line represents best fitting linear model, hatched lines represent 95% confidence interval. r-values in italics represent results for GABA scaled to total Creatine.

BOLD may influence measurements of GABA by masking the available signal from proton-MRS measurements. We assessed the impact of BOLD-effects on GABA, by correlating the line width of the total Creatine singlet at 3.03 ppm, a general measure of MRS signal quality, with GABA values. There was no correlation (Fig. 5B, GABA:H_2_0, Pearson’s Linear Correlation, r = 0.34, p =0.1, GABA:tCr, Pearson’s Linear Correlation, r = 0.15, p = 0.47). On the other hand, we can demonstrate that GABA estimates correlate with reliability of GABA fit (GABA:H_2_0, Spearman’s Rank Correlation, r = -0.74, p < 0.001; GABA:tCr, Spearman’s Rank Correlation, r = -0.90, p < 0.001), and further that GABA and glutamate during visual stimulation were highly correlated (metabolite:H_2_0, Pearson’s Correlation, r = 0.43, p = 0.03; metabolite:tCr, Spearman’s Rank Correlation, r = 0.491, p = 0.016). These findings rule out the possibility that spectral quality difference and BOLD effects systematically affected overall GABA estimates.

## DISCUSSION

Eye dominance in the normal visual system may be regarded as a window into how the brain selects and combines information from the two eyes. Extreme eye dominance has a pathological form manifest in ‘amblyopia’, which impairs normal visual function. We demonstrate a link between intracortical inhibition and normal eye dominance in the healthy visual system, based on a novel approach to measurement of inhibition in the cortex. We applied monocular visual stimulation during combined fMRI-MRS at 7-Tesla. There was no link between eye dominance and inhibition in the simultaneously acquired BOLD-signal, or in resting GABA levels.

We show that GABA-levels measured during visual stimulation are linked to eye dominance in the healthy human brain. Our results are in agreement with the view that the neural mechanism of eye dominance is mediated by GABAergic inhibition [9, 28]. In regard to the difference in GABA between DE and NDE eye (‘interocular GABA’), individuals with stronger eye dominance showed a greater difference in GABA. Eye dominance may therefore be due to a systematically weaker ability of the non-dominant eye to inhibit the stronger eye during active viewing. In confirmation, no relationship to eye dominance was present when GABA levels were measured during the resting state, during which participants had their eyes closed.

What might the visually driven GABA signals represent? We note that our measures of GABA levels were taken during continuous visual stimulation, engaging attentional mechanisms and low-level sensory processing. Given that MRS-visible GABA is unlikely to comprise a direct reflection of synaptic GABA release, we speculate that the MRS-visible signal may represent changes in tonic inhibition following neural activation. While the relationship between neuronal activity and ambient GABA levels in the visual cortex is still being investigated, it is known that synaptic release of neurotransmitter can spill over from the synaptic cleft into the extrasynaptic space [29]. This can cause changes in the ambient GABAergic tone [30], currently thought to be the main source of the MRS visible GABA signal [21, 31, 32]. Our findings here emphasize that revealing of inhibitory interactions in the binocular visual system depends on the status of visual input.

There was no relationship between eye dominance and the interocular BOLD-signal. Only a handful of studies have investigated this relationship and it is fair to say that the results have so far been inconclusive, partly due to differences in how eye dominance was assigned. Greater BOLD-activation in V1 has been found during DE versus NDE stimulation [33-35], whereas others find no difference [36]. In those studies, eye dominance was assigned using ‘sighting dominance’, which evaluates which eye is used for viewing a distant target. Different methods of assigning eye dominance often do not agree with one another (for a review see [37]). We measured sighting dominance for all our subjects and found that it corresponded with assignments of sensory eye dominance assignments in only 7 out of 12 participants.

A better comparison with our study is found from a recent study using binocular rivalry: here they reported that increases in eye dominance induced by monocular deprivation are linked to increases in deprived eye BOLD-activation [38]. The lack of any correlation in our BOLD-signal data are consistent with the view that the BOLD-signal cannot detect inhibitory signals [39], whereas MRS can peform this to a significant degree [17, 18, 20].

There are other limitations to our study. Firstly, our study has a relatively small sample size (N = 12). However, our methods replicate a recent, previous result [18], performed with a greater number of participants (N = 21), different field strength (3T) and MRS sequence. This suggests that our sample size and data quality contained sufficient power to replicate previous findings for GABA. Secondly, our MRS-voxel size did not permit the sub-millimeter resolution necessary to give direct imaging of ocular dominance columns, within which lateral inhibition is thought to drive eye dominance. Whilst we are confident that measured GABA concentrations are specific to dominant or non-dominant eye stimulation, a long-term improvement in the spatial resolution of MRS imaging is required to bring the imaging of ocular dominance columns within reach.

In conclusion, we discover that the healthy human visual cortex displays a relationship between the perceptual measure of eye dominance and the neural measure of GABAergic inhibition. This relationship is specific to the conditions of visual stimulation, but confirms and extends a long-standing hypothesis that imbalances in cortical inhibition during monocular visual response are related to eye dominance [9, 15, 16]. The extent to which these relationships can be related to clinical conditions of extreme eye dominance are being currently investigated in our laboratory.

## SUPPLEMENTARY MATERIALS

### Individual MR-Spectra

**S1.**
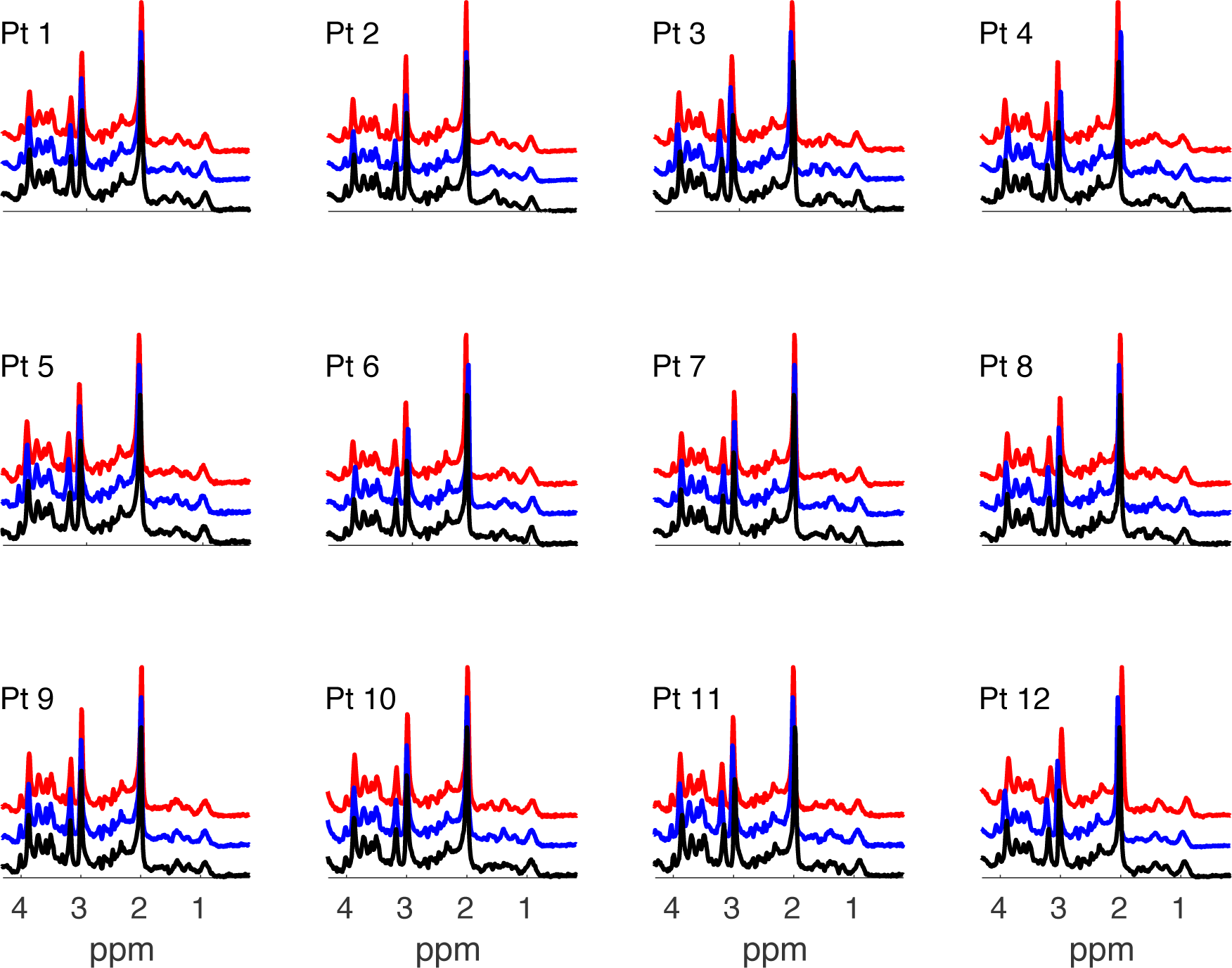
Raw MR-spectra for all 12 participants and all conditions (red, dominant eye stimulation, blue, non-dominant eye stimulation, black, rest) normalised to the total Creatine singlet at 3.03 ppm. X-axis denotes chemical shift axis in parts per million (ppm).

## METHODS

### Participants

Thirteen participants (7 females, 29.2 ± 6.0 years) including two of the authors took part in the study. All data presented were collected as part of a behavioural and MRI-data set, of which a subset has been published [24]. One participant was identified as an outlier due to a large percentage (68.5%) of mixed periods in the binocular rivalry experiment. Participants had normal or corrected-to-normal vision, no neurological impairments and normal stereo-acuity (< 120 arc sec, TNO Stereo test, Lameris, Utrecht). All subjects were involved in a 1-h psychophysical session to measure binocular and monocular visual function, as well as eye dominance, and took part in a 1.5-h MRI session to measure interleaved changes in neurochemical concentrations and BOLD-activity. All subjects gave written informed consent. Approval for the study was obtained by the University of Oxford Research Ethics Committee (MSD-IDREC-C1-2014-146). The study was carried out in accordance to the Declaration of Helsinki.

### Behavioural protocol and procedure

Participants’ eye dominance was measured in a separate psychophysical session outside of the scanner (Fig. 1A). While both sighting and sensory eye dominance tests were collected, only sensory eye dominance was used to compare to imaging data as it quantified the strength of eye dominance. The Miles Test [40] was used to measure sighting eye-dominance, the dominant eye was the eye used for sighting of a distant target through an aperture. Binocular rivalry was used to obtain a measure of sensory eye dominance. Head position was stabilised with a chin and headrest. Stimuli were displayed on two gamma-linearised CRT monitors (viewing distance 57 cm) viewed through a Wheatstone mirror stereoscope. Stimuli were generated using MATLAB (The MathWorks, Natick, MA) with Psychophysics Toolbox [41] running on a mac-mini. Binocular rivalry stimuli were two achromatic gratings (orientation: ±45 deg off vertical, spatial frequency: 6 cycle/deg, contrast: 100%, diameter: 3.2 deg) presented on a uniform mid-grey background in the centre of vision. After successfully fusing a Nonius fusion target, participants self-initiated the trial and reported the perceived orientation of a tilted grating using the left (counter-clockwise) and right (clock wise) arrows of a computer keyboard, and pressed the upward arrow when a mixed percept was perceived. The orientation of the grating was randomised across runs between the left and right eye. After a practice run, all participants took part in three binocular rivalry runs each lasting 180 s.

### Binocular rivalry analysis

Data were first pre-processed: missing data before perceptual transitions were assigned to the subsequent percept. Responses of < 200 ms, mixed responses or missing data with no response were removed from the analysis. Because of the skewedness of phase durations (Fig. 1B), the median phase durations were calculated [27]. To quantify sensory eye dominance, an ‘eye dominance index’ (EDI) was calculated as:

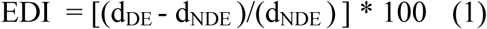

Where d_DE_ is the median phase duration through the dominant eye and d_NDE_ is the median phase duration through the non-dominant eye. The EDI quantifies the percentage increase in median phase duration of the dominant over the non-dominant eye.

### Visual stimulation in MRI scanner

The visual stimulation was designed to continuously engage visual processing (Fig 2A). The same visual stimulation was viewed with the left eye or the right eye. The non-viewing eye was occluded with a translucent, form-depriving occluder [20]. In order to minimise head motion, participants practised switching the occluder to the non-viewing eye prior to scanning and were encouraged not to give any verbal responses while in the scanner. The neurochemical concentrations were measured as total concentration across scans, as the continuous fixation task at the centre of the display provided visual stimulation and attentional deployment [42] for interocular suppression to be engaged. As a control analysis, we also performed the neurochemical analysis using the difference between checkerboard > baseline.

Stimuli were generated on a Mac Book Pro workstation using Matlab and Psychtoolbox-3 [41] and displayed on a gamma-linearised Eiki LC-XL100 projector (resolution: 1024 × 768, refresh rate: 60 Hz). Participants viewed the stimuli through 45° angled mirrors on a back-projection screen (viewing distance: 60 cm). As shown in Fig 1B, visual stimuli were full-field checkerboards, contrast reversing at 8 Hz (stimulus size = 19.82° x 14.25°, 8 Hz flicker, mean luminance=385 cd/m^2^; 50% contrast). The baseline condition was a uniform black screen (2.33 cd/m^2^) with a fixation dot. Each run consisted of four alternations of baseline (64 s) followed by flashing checkerboards (64 s). A central fixation dot was displayed (white with black border, size = 0.75°) throughout the experiment. Participants were instructed to maintain fixation and press any button on a MRI-safe button box when the marker turned red (500 ms, ∼once in every three seconds).

### MR protocol

Magnetic Resonance data were acquired using a 7T whole body (Siemens, Erlangen) MR-scanner with a Nova Medical head coil (single transmit, 32 receive channels). A T1-weighted structural scan was acquired for each participant with a 1-mm isotropic resolution (MPRAGE, repetition time TR = 2.2 s, inversion time T_I_ = 1.05 s, echo time TE = 2.82 ms, FOV= 192 × 192 x 176 mm, flip angle = 7°, total acquisition time = 171 s) to permit placement of the visual cortex voxel-of-interest (VOI). A 2 × 2 x 2 cm (8 cm^3^) MRS VOI was positioned in the occipital lobe, centred on the calcarine sulcus and at the midline to include V1 in both hemispheres. fMRI and MRS data were acquired using a combined fMRI-MRS sequence (Ip et al., 2017), yielding a slab of EPI (3D EPI, resolution= 4.3 × 4.3 × 4.3 mm; flip angle = 5°, repetition time TR_epi_= 40 ms, TE= 25 ms, FOV= 240 mm, 16 slices) and a MR spectrum upon each TR. For each image experimental run, 128 spectral averages were acquired using short-echo semi-localisation by adiabatic selective refocusing (semi-LASER) pulse sequence (TE = 36 ms, TR_mrs_ = 4 s) with VAPOR water suppression and outer volume suppression [43, 44]. Data were collected as single free induction decay per excitation. A dielectric pad measuring 110 × 110 x 5 mm^3^ containing a suspension of Barium Titanate (BaTiO_3)_ and deuterated water (mass-mass ratio of 3:1) was placed behind the occiput of each participant [45, 46] to increase the effective transmit field efficiency close to the pad [47].

### fMRI analysis

fMRI data were analysed using FEAT (FMRI Expert Analysis Tool) v. 6.00, part of the FSL software distribution (FMRIB’s Software Library, www.fmrib.ox.ac.uk/fsl; RRID:SCR_002823). Pre-processing included motion correction MCFLIRT [48]; non-brain tissue extraction [49]; 5 mm smoothing, grand-mean intensity normalization and high pass temporal filtering (Gaussian-weighted least squares straight line fitting, main experiment = 132 s). Registration of EPI to an initial 2-mm structural image used 6 DOF, followed by registration to the 1-mm isotropic T1-weighted structural image using boundary-based registration (BBR) in FLIRT [48, 50]. BOLD-change in the MRS-voxel was calculated using Featquery.

### fMRS analysis

Pre-processing for raw MRS data was performed in MRspa (https://www.cmrr.umn.edu/downloads/mrspa/) and included eddy current correction, frequency alignment to the tNAA singlet at 2.01 ppm and phase correction using a least-square algorithm. Metabolites within the chemical shift range of 0.5 to 4.5 were analysed using LCModel [51, 52]. Group responses were the average across individual participant values (Fig S1). Metabolite levels were estimated using a basis set of alanine (Ala); ascorbate/vitamin C (Asc); aspartate (Asp); glycerophosphorylcholine (GPC); phosphorylcholine (PCho); creatine (Cr); phosphocreatine (PCr); γ-aminobutyricacid (GABA); glucose (Glc); glutamine (Gln); glutamate (Glu); glutathione (GSH); inositol (Ins); lactate (Lac); phosphoeth anolamine (PE); scyllo-inositol (sIns); taurine (Tau); N-acetyl-aspartate multiplet (mNAA); N-acetyl-aspartate singlet (sNAA); Acetyl moiety of N-acetylaspartylglutamate (sNAAG); aspartyl moiety of NAAG (mNAAG); glutamate moiety of NAAG (gNAAG). Macromolecule inclusion procedures were performed as in Bednarik et al., (2015).

The percentage of cerebro-spinal fluid (CSF, 8.00 ± 3.05%), white matter (WM, 48.83 ± 4.82 %) and grey matter (GM, 43.16 ± 3.59%) in the V1 MRS voxel from the T1-weighted high resolution anatomy scan was determined using automated tissue segmentation (FSL v6.0 FAST [53]) and custom written scripts. We identified the impact of BOLD-effects in metabolite spectra by estimating the total Creatine line width at 3.03 ppm (tCrLW) during dominant and non-dominant eye runs [54]. Metabolites were quantified relative to the same unsuppressed water spectrum, collected at the beginning of each MRI-session. Water referenced metabolite values were corrected for the amount of CSF using the equation : [M_corr]_ = [M] x (1 / [1 – f_CSF_]), where M_corr_ is the corrected metabolite concentration, M the metabolite value from LCModel and f_CSF_ the CSF fraction [20, 55]. To ensure that the main results were not driven by the reference method, metabolite:tCr correlations were also used and presented alongside results from metabolite:H_2_0 values. Metabolite ratios were corrected for proportion of gray matter and white matter fractions using equation (2) from Harris et al., 2015:

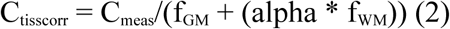

where c_tisscorr_ is the corrected metabolite ratio, c_meas_ is the uncorrected metabolite ratio, f_GM_ and f_WM_ are the participant’s GM and WM fractions, and α is the ratio of the metabolite concentration in grey and white matter, set to 0.5.

The reliability of the MRS fit were quantified by the Cramer-Rao lower lounds (CRLB), and the criterion of 30% across participants (128 spectra/participant, N = 12) chosen to exclude poorly modelled metabolites. The averaged tissue corrected metabolite levels across 12 participants (128 spectra/participant) for GABA:H_2_0 was 1.30 ± 0.14 (GABA:tCr, 1.19 ± 0.13, CRLB = 28.23±3.27 %). We also calculated the coefficient of variation (CoV) between repeated measurements of the estimated signal, to reflect the ability to detect physiological signals independent of the noise [43]. If the CoV is higher than CRLB, it means that the signal is indistinguishable from the noise. If CoV is equal to or lower than CRLB, relatively small variations in signal may be measurable. To estimate within-subject CoV, for each condition and subject, data were split into the first (64 spectra) and second half (64 spectra) of the scan, and analysed using LCModel. The CoV was calculated by dividing the standard deviation of the two measurements by their mean concentration [56]. Average GABA concentrations from the first and second half were highly correlated (Spearman’s Rank Correlation, R = 0.834, p = 0.001). The within-subject CoV was 8.88 ± 3.10 STD, in the range of previously reported within-subject CoV for GABA using non-edited MRS sequences [8-12%] [57, 58].

The interocular difference metric in neural response is calculated for MRS measured GABA and for the %BOLD-signal change using the same equation:

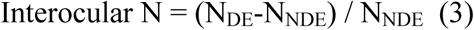

where N_DE_ is the neural response value during dominant eye viewing, N_NDE_ is the neural response value during non-dominant eye viewing.

### Statistical Analysis

Data were assessed for normality using the Shapiro-Wilk Test. If normality was refuted, (SW-test, p < 0.05) non-parametric statistics were used. The Wilcoxon’s Signed Rank test was used to test for significant differences in the median between two repeated measures. Spearman’s Rank correlation coefficients were calculated to evaluate relationship between two independent variables of interest. The p-value was calculated using the exact distribution test. If normality was not refuted, paired t-tests were applied to test for differences between group means. Pearson’s Linear Correlation coefficients were computed to assess the relationship between two independent variables. In all cases, the significance value alpha was set to 0.05, uncorrected. Due to the low number of participants, it is possible that extreme data points influenced the data. However, no outliers were identified in metabolite or eye dominance measures using the function boxplot.m in Matlab.

## Acknowledgements

This work was supported by the Medical Research Council (MR/K014382/1), The Royal Society (University Research Fellowship to HB), and the French National Research Agency (ANR-19-CE28-0008, PlaStiC). We would like to thank Dr William Clarke for advice on MRS quantification. The Wellcome Centre for Integrative Neuroimaging is supported by core funding from the Wellcome Trust (203139/Z/16/Z).

## Author contributions

Conceptualization, I.B.I., C.L., H.B.; Methodology, I.B.I., U.E.E., C.L., H.B.; Investigation, I.B.I. U.E.E.; Writing, I.B.I; Review and editing, I.B.I, U.E.E., C.L., H.B., A.J.P.; Funding Acquisition, A.J.P., H.B.; Resources, C.L., U.E.E.; Supervision, H.B.

